# A three-photon head-mounted microscope for imaging all layers of visual cortex in freely moving mice

**DOI:** 10.1101/2022.04.21.489051

**Authors:** Alexandr Klioutchnikov, Damian J. Wallace, Juergen Sawinski, Kay-Michael Voit, Yvonne Groemping, Jason N. D. Kerr

## Abstract

Recent advances in head-mounted microscopes have enabled imaging of neuronal activity using genetic-tools in freely moving mice but these microscopes are restricted to recording in minimally lit arenas and imaging upper cortical layers. Here we built a 2 gram, three-photon excitation-based microscope, containing a z-drive that enabled access to all cortical layers while mice freely behaved in a fully lit environment. We show that neuronal population activity in cortical layer-4 and layer-6 was differentially modulated by lit and dark conditions during free exploration.

---

Recently developed multiphoton head-mounted microscopes^1–4^ can utilize, with single-cell resolution, the vast range of genetically encoded tools^5^ used for recording neuronal activity in freely behaving mice. Two-photon excitation-based head-mounted microscopes are restricted to imaging neuronal activity in upper cortical layers^1,2,4^, whereas three-photon excitation (3PE)-based headmounted microscopes, while removing this imaging depth limitation^6^, have so far been too physically restrictive to take full advantage of mouse-based molecular tools. To gain access to larger numbers of neurons and remove the need to interfere with an animal’s behavior when changing the focal plane, remote focusing has been applied to several two-photon excitation-based head mounted microscopes^2,4^. While this has increased focusing range by several hundred microns and allowed volume imaging at low frame rates, acquisition is still limited to the upper cortical layers and the microscope weight has markedly increased^2^. Lastly, as mice actively sense their environment using vision^7,8^, utilizing their full behavioral repertoire requires a fully lit visual environment, which is problematic for head mounted microscopes using photon multiplier tubes due to their sensitivity^1,2,4,6,9^. Ideally, a head-mounted multiphoton microscope could image from all cortical layers, be shifted at will between layers and be usable in a fully lit environment to allow the animals’ to access their full sensory repertoire^7,8^.

We designed a head-mounted three-photon excitation microscope^6^ specifically for imaging activity from all cortical layers in mice. Our new optical pathway was optimized to allow remote focusing^10^, gaining access to all cortical layers (Fig. 1a&b) and contained a modified detector system to allow imaging in a lit arena. A remote focusing mechanism^10^, with a sufficient range to cover the entire cortical depth, required an optical pathway that maintained diffraction limited imaging across the entire focus range. Like with previous bench-top designs^11^ we incorporated a z-drive that changed the collimation properties (defocus) of the beam, resulting in changes in the working distance (Fig. 1c). The resulting optical system had a simulated working distance ranging from ~1.1 to 1.9 mm, sufficient for moving the imaging depth from cortical layer 1 to the corpus callosum in the mouse (~800 μm, Fig. 1a). The optical simulation predicted that imaging was diffraction limited over almost the entire range (> 700 μm in z with a FOV > 400 μm, Supplementary Fig. 1), while still maintaining a sufficient excitation numerical aperture (NA=0.4) for sub-cellular resolution^12^. Although higher excitation NA was possible, there was a trade-off between achievable z-range and resolution (Supplementary Fig. 2). To remotely change the working distance during free behavior, removing the need to interfere with the animal, we built two separate defocusing mechanisms that both could utilize the microscopes optics to change the working distance (Fig. 1c, Supplementary Fig. 3). Using the maximum tuning range of an electrically tunable lens^13^, the image plane could be moved over a range of ~400 μm while maintaining a resolution required for sub-cellular imaging (axial resolution < 17 μm full width half maximum (FWHM), Supplementary Fig. 3). While this lens provided fast access to any imaging plane within the 400 μm range, and potential for multiplane imaging^4^, the electrically tunable lens increased the total microscope weight by ~40% (> 800 mg). To further reduce the weight while enabling access to the entire cortex, we designed a z-drive based on a lightweight mechanical system capable of moving the imaging plane from cortical layer 1 to the corpus callosum (Fig. 1c). The design exploited the beams divergence when exiting the fiber-tip, enabling changes in the working distance as the distance between the fiber-tip and the collimation lens changed (Fig. 1c). A 12 mm movement combined with a collimation lens with a 6 mm effective focal length (EFL) produced a >700 μm z-range (Fig 1d). With this optical configuration, the field of view was > 300 μm (Fig. 1b, limited by the scanner specifications), with subcellular lateral and axial resolution being maintained (< 1.2 μm lateral and < 20 μm axial, both FWHM, Fig. 1e). This approach added minimal additional weight (< 100 mg), and the smaller specified EFL of the objective lens (1.7 mm vs 3 mm in previous design^6^ reduced the weight of the optical system compared to our previous miniature 3PE microscope (720 mg vs >2g in previous design^6^). Together these optimizations resulted in a total microscope weight of 2 g. The resulting miniature microscope provides access to the full cortical thickness (Supplementary movie 1) in a freely-moving mouse without the need to interfere with the animal’s behavior. To confirm the range and stability of the microscope and z-drive mechanism, we created a mouse line expressing Cre recombinase both in layer 4 (L4)^14^ and 6 (L6)^15^. Using an AAV encoding cre-dependent jGCaMP7f we could then drive indicator expression selectively in those cortical layers (Fig. 1f). In single imaging sessions we could sequentially measure calcium transients from neuronal populations in L4 and L6 (Fig. 1g&h, N=9 L4 populations from 3 animals, N=9 L6 populations from 4 animals), or measure from multiple image planes within a layer (Fig. 1i), and could make repeated measurements from these cortical layers in sessions spread over multiple days (N=9 sessions total, from 4 animals, number of imaging days 2.3 ± 1.3, mean ± SD, range 1 to 4, average days post window implant 5.7 ±2.7 days, mean ± SD, range 2 to 10). The microscope configuration described here provided a field of view of ~300 μm diameter, capturing in these experiments an average of 34.2 ± 15.7 neurons per FOV for imaging in L4 (mean ± SD, range 20 to 60, N=9 FOVs) and 98.7 ± 25.8 for L6 (mean ± SD, range 68 to 141, N=9 FOVs). During these imaging sessions, animals were freely exploring a linear track (96.0 x 9.7 cm, L x W), and were mobile and active, with average movement path length per imaging file of 661.9 ± 299.3 cm (mean ± SD, range 142.3 to 1582.7 cm, n=30 imaging files from 18 populations), with an average velocity (while moving) of 9.0 ± 0.85 cm/s (mean ± SD, range 7.8 to 10.5 cm/s, n=30 imaging files from 18 populations). Overall, imaging was stable (Supplementary movie 2), and data excluded for all reasons (including entanglement of the optical fiber and electronic cables, image brightness fluctuation due to shifting of the immersion solution and excessive image motion) was for the majority of imaging files (20/31 files) <3% of the recorded data (average data rejection for all files 10.3 ± 17.6%, mean ± SD, range 0 to 72.1%, N=31 imaging files).

**Figure 1.**
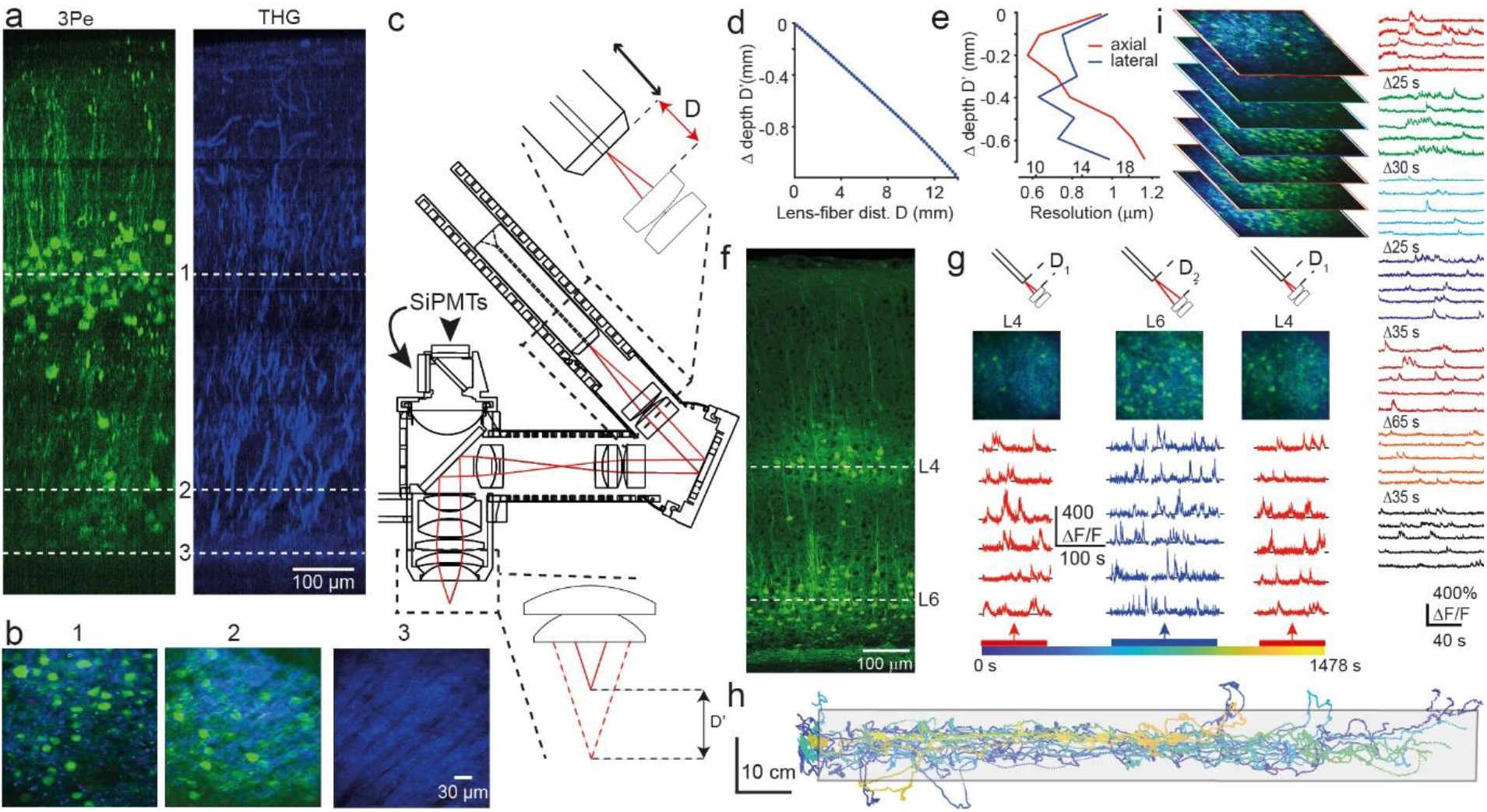
A light-weight miniature 3-photon fiberscope with z-drive for imaging in freely moving mice. **a**, image stack in side-projection showing jGCaMP7f-labelled neurons (left) and third harmonic generation signal (right) acquired with the microscope mounted on an external micromanipulator. Dashed lines indicate the image planes for the corresponding images shown in b. **b**, individual images from the data shown in a. **c**, microscope schematic showing the mechanism used to shift the image plane. Shifting the fiber tip through distance D (top insert) results in a shift in the image plane of D’ (bottom insert). **d**, zemax-simulation of relationship between the lens-fiber-tip distance (distance D in c) and corresponding image plane depth (distance D’ in c). **e**, dependence of axial (red) and lateral (blue) resolution on image plane depth, measured in a sample of 0.5 mm fluorescent beads. Inner x-axis scale refers to axial- and outer scale to lateral resolution. **f**, histological section showing jGCaMP7f labelling in layer 4 and 6 of mouse primary visual cortex. **g**, overview images (middle) and example Ca^2+^-fluorescence traces (lower) from sequential imaging of neuronal populations in layer 4 (red) and 6 (blue) in the same behavioral session from one animal. Color-coding in the time scale (bottom) corresponds with the color-coded position of the animal on the linear track shown in h. Top schematics illustrate the corresponding relative fiber-tip to lens distances (dist. D in c). **h**, animal position on the linear track during the behavioral session in which the data in h were acquired. Color-coding corresponds with the color-coded time scale in g. **i**, overview images and example Ca^2+^-fluorescence traces from sequential imaging of multiple planes of labelled neurons in layer 6 acquired in one behavioral session (Δ denotes time from end of previous plane until start of the new plane). Colored border on overview images corresponds to the color of the example Ca^2+^ traces. Times under the colored blocks of Ca^2+^ traces indicate acquisition start times for the corresponding image plane in the behavioral session. Scale below bottom traces apply to all Ca^2+^ traces.

To reduce the constraints on the animal’s behavior^3^ of previous miniature microscope designs^1,2,4,6^ and simultaneously to increase light-collection efficiency, we removed the stiff plastic optical fiber (POF, NA = 0.63, 1mm diameter) and designed a 2-channel detector system that was mounted on the microscope (Fig. 2a, Supplementary Fig. 4). The detectors were silicon photomultipliers (SiPMTs), which are compact and lightweight (2*3 mm^2^, 5 mg) with a large detection area (1.3*1.3 mm^2^) and high detection efficiency (> 60%) and gain (> 10^6^). Under matched imaging conditions, the SiPMTs achieved a higher signal-to-noise ratio than the POF detection system (factor 1.8 higher, Fig. 2b) over a larger FOV (Supplementary Fig. 4). Unlike GaAs-PMTs^6,9,16^ SiPMTs were resilient to stray light-exposure. To reduce the high level of dark counts (~10^4^ counts*mm^-1^s^-1^) generated by these SiPMTs we implemented dark count rejection, by integrating the fluorescence signal in a short time window after each laser pulse^6^. This approach collected only emitted fluorescence generated by the excitation pulses (Fig. 2c, max 1MHz excitation pulse rate, 50 ns integration window, rejection ratio 20 times for 1 MHz pulse rate). To take advantage of the SiPMT detection system for visually based experiments, we designed a system that timed the experimental arena’s lighting to switch on only during the interval between acquisition of each line (3 kHz) that forms the full-frame (Fig. 2d), a flicker rate well above what can be detected by rodents^17,18^. This allowed imaging of the same neuronal populations during either fully lit or darkened conditions (Fig 2d-e) without significantly changing the detected fluorescence in the image (Fig 2f left, average pixel intensity over the full frame one frame before and after light-dark transition, N=22 transitions from 3 animals, P = 1, two-sample Kolmogorov-Smirnov test). Further, Ca^2+^ fluorescence traces recorded from labelled structures imaged across a transition from light to darkness or darkness to light showed no artefacts in the fluorescence traces (Fig. 2f right, Supplementary fig. 5, Supplementary Movie 3), apart from a transient re-synchronization artefact which occurred in some cases on activation of the environmental lighting (mean ± SD number of frames with artefacts 1.9 ± 4.8, range 0 to 21, N=20 dark to light transitions from 3 animals).

**Figure 2.**
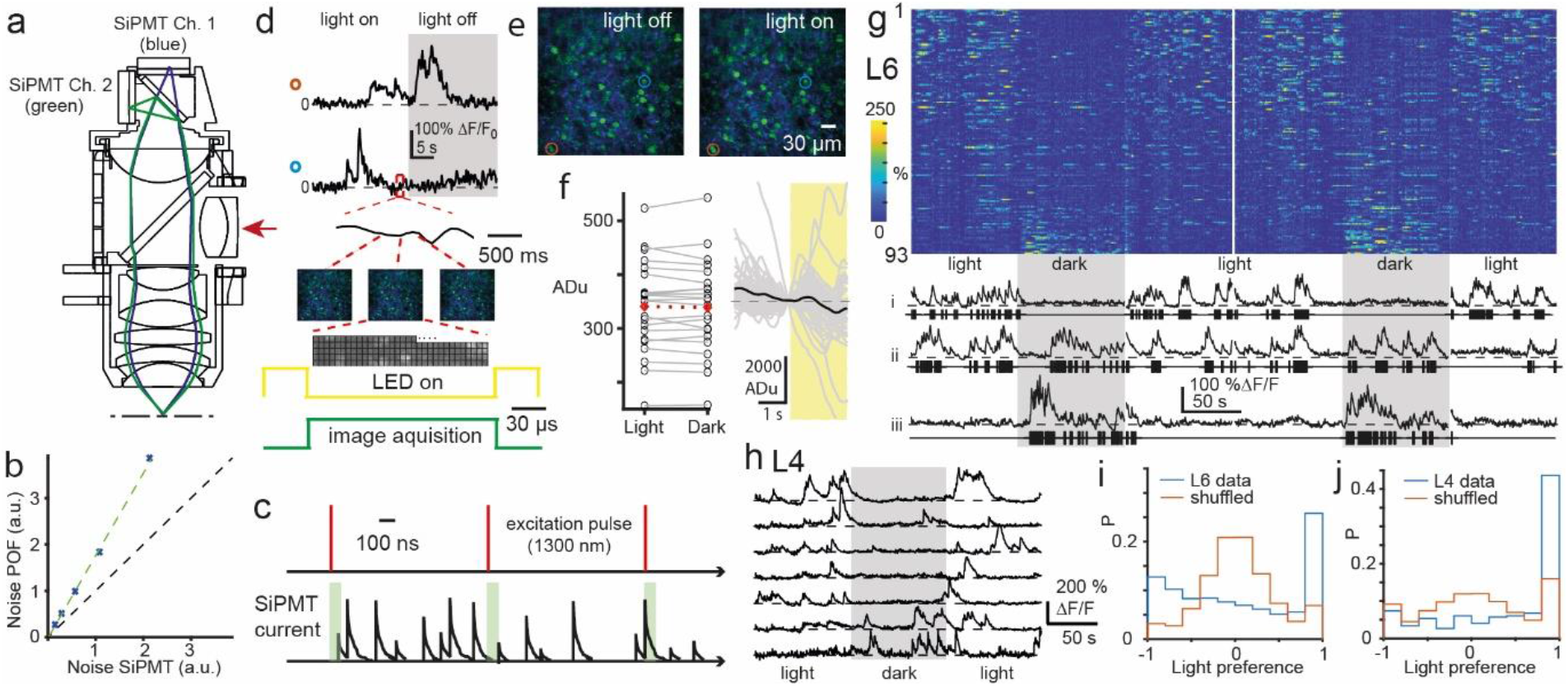
Adaptions allowing imaging in a lit environment. **a**, microscope schematic highlighting the detector system, showing light paths and on-board detectors for green (green) and third harmonic generation signal (blue) channels. Scanner and z-drive optics omitted. Direction of excitation path indicated by red arrow. **b**, quantification of image noise using on-board detectors (SiPMT) and remote PMTs coupled via a plastic optical fiber (POF). **c**, schematic of pixel-wise timing of excitation pulses (upper) and emitted fluorescence integration window (lower, green boxes) used for dark count rejection. **d**, example Ca^2+^-fluorescence traces from the neurons indicated in e during a transition from the environment being lit to being dark, with schematic showing timing of line-wise image acquisition and environmental lighting activation. The short segment of the lower trace outlined by the dashed red box is shown expanded for illustration. Three consecutive full frame images are shown to illustrate the activation of the environmental lighting (yellow) between acquisition of image lines (green). **e**, example images (average of 50 frames) acquired with environmental lighting off (left) or on (right). **f**, average pixel intensity over the whole frame from green channel data one frame before (Light) and one frame after (Dark) transitions from light to dark. Open circles show data from individual transitions, red diamond represents the mean (N=22 transitions, p=1, two-sample Kolmogorov-Smirnov test). **g**, color-coded Ca^2+^ fluorescence traces for all neurons in an example L6 population (upper) and example fluorescence traces from three neurons (lower), in two consecutively acquired imaging files. Both imaging files have transitions between environmental lighting being on (light) and off (dark, grey boxes). Color-coded fluorescence traces are sorted by neuronal light-dark preference index. Black ticks below example Ca^2+^ traces indicate inferred action potential firing. Time scale in the individual Ca^2+^ traces applies to both the color-coded plot and the individual traces. Fluorescence scale bar applies to all individual Ca^2+^-traces. **h**, example Ca2^+^-fluorescence traces from a L4 population with periods where the environmental lighting is on or off (gray box). **i**, distributions of light-dark preference index (blue) for all recorded neurons in L6 in periods where the animal was mobile (vel. > 5cm/s), including also distributions for shuffle-controlled firing rates (red, N=880 neurons from 9 datasets from 4 animals). j, as for I, but for all neurons in L4 (N=298 neurons from 9 datasets from 3 animals).

As previous work has suggested that neuronal activity in L6 and L4 is differently modulated in dark and light conditions in head restrained and freely moving mice^19,20^, we next compared neuronal activity from populations of either L4 or L6 neurons while mice were freely exploring the linear track in either lit (35 lm/m^2^) or dark (~0 lm/m^2^) conditions (Fig. 2g, Supplementary Movie 3). The mice were active in both conditions (total path length lit 231.6 ± 169.9 cm, dark 276.7 ± 153.9 cm, mean ± SD, p=0.12, Wilcoxon rank sum test, N=18 datasets from 4 animals), though, when moving, moved significantly slower in the dark (velocity when moving lit 9.26 ± 1.09 cm/s, dark 8.49 ± 0.98 cm/s, p=3.5*10^-2^, Wilcoxon rank sum test, N=18 datasets from 4 animals). Across L6 populations, neuronal activity ranged from active neurons in lit conditions becoming almost silent upon transition to dark (Fig. 2g trace i) to neurons that had sparse activity during lit conditions becoming active in darkened conditions (Fig. 2g trace iii). In contrast, L4 population activity was elevated in the light and sparse in the dark, but few if any neurons with elevated activity were observed in darkness (Fig. 2h). To facilitate statistical comparisons of neuronal activity under these different conditions we inferred neuronal spiking from the Ca^2+^ fluorescence traces using a previously published algorithm^21^ (see inferred activity in Fig. 2g, traces i-iii and Supplementary Fig. 6), and used the inferred spikes to calculate a light preference index (LPI, see Methods). As neuronal activity in V1 can be modulated (either enhanced or suppressed) by vestibular signals in lit or dark environments differentially^19^, we split our datasets into epochs where the animal was mobile or stationary, and consequently with greater or lesser vestibular signal respectively. When the animal was active, significant populations of neurons whose activity was strongly enhanced either in light (LPI around 1) or dark (LPI around - 1) were observed in L6 (Fig. 2i, observed distribution of LPIs vs LPIs from shuffled data p=4.3*10^-36^, two-sample Kolmogorov-Smirnov test, N=9 datasets from 4 animals). A significantly greater proportion of the recorded neurons fell into the extreme LPI categories when the animal was moving compared to when stationary (L6 LPI distribution stationary vs. mobile, p=1.9*10^-5^, two-sample Kolmogorov-Smirnov test, N=9 datasets from 4 animals, Supplementary Fig. 7), consistent with modulation of neuronal activity by motion driven vestibular input^19^. In L4 populations, while neurons with activity enhanced by light were observed (Fig. 2j, observed distribution of LPIs vs LPIs from shuffled data p=1.5*10^-26^, two-sample Kolmogorov-Smirnov test, N=9 datasets from 3 animals), the distribution representing neurons with activity enhanced in darkness (LPI around −1) was not different to that observed in shuffled data (Fig. 2j). Here we present a miniature headmounted three-photon microscope with remote focusing, suitable for imaging activity from neuronal somata and dendrites from all cortical layers in freely moving mice. This microscope contained an onboard 2-channel detector system that was robust to ambient light, and a scanner-synchronized arena lighting system which together enabled imaging of neuronal activity in a fully lit behavioral arena. This microscope allows the full suite of molecular tools and mouse lines to be brought to bear to study cortical function from any depth in the cortical mantle of a freely moving and behaving mouse.

## Materials and Methods

### Animals and surgical procedures

All animal experiments were conducted in accordance with institutional animal welfare guidelines and with animal experimentation approval granted by the Landesamt für Natur, Umwelt und Verbraucherschutz Nordrhein-Westfalen, Germany.

Mice were housed in an SPF temperature- and humidity-controlled facility on a 12 h light–dark cycle with food and water available ad libitum. Mice were group housed until surgery and singly housed afterwards. Mice (2 males, 44 females) were 8–14 weeks old (average of 11 weeks) and weighed 20–24 g (average of 22 g) at the time of the virus injection.

For this study we used mice that resulted from crossing heterozygous animals from two Cre-expressing lines, Ntsr1-Cre mice and Scnn1a-Cre mice. *Ntsr1* (neurotensin receptor 1)-Cre mice (B6.FVB(Cg)-Tg(Ntsr1-cre)Gn220Gsat/Mmcd) were obtained from the Mutant Mouse Resource and Research Center (MMRRC, #030648-UCD) and donated by Nathaniel Heintz and Charles Gerfen, with Cre recombinase predominantly expressed in Layer 6 of the cortex^15^. *Scnn1a* (sodium channel, nonvoltage-gated 1 alpha)-*Cre* mice (Tg(Scnn1a-cre)3AibsTg(Scnn1a-cre)3Aibs) were obtained from the Jackson Laboratory (#009613) and donated by Ed Lein and Theresa Zwingman with Cre expression in cortex layer 4^14^. The Cre-animals were maintained in a heterozygous state with C57BL/6J background. Expression patterns of these transgenic Cre lines have been described in the original papers cited.

The presence of the individual transgenes was confirmed by genotyping PCR with layer-specific primers. Primers for Snn1a-Cre animal specific PCR were 5’-AAAGAGAAGCGGGAGTCAG-3’ and 5’-GACCGGCAAACGGACAGAAG-3’. Primers for genotyping of Ntsr1-Cre animals were 5’-TCCCAGGATCTCCTGGATAG-3’ and 5’-GACCGGCAAACGGACAGAAG-3’ (forward and reverse primer, respectively).

For virus injections AAV1/2.hSyn.FLEX.jGCAMP7f was purchased from Addgene, the plasmid pGP-AAV-syn-FLEX-jGCaMP7f-WPRE was a gift from Douglas Kim & GENIE Project (Addgene plasmid #104492)^22^.

#### Genotyping PCR

Genomic DNA was isolated from ear biopsies using lysis buffer (10 mM Tris/HCl pH 8.0; 100 mM EDTA pH 8.0; 0.5% SDS) containing 50 μl Proteinase K [10 mg/ml] with subsequent isoropanol precipitation and ethanol wash step. The polymerase chain reaction (PCR) was performed using DreamTaq Master Mix (ThermoFisher) and construct specific primers.

#### Surgical implants and mounting devices

The equipment used to mount the microscope on the head of the animal for freely moving experiments consisted of a surgically implanted headplate, an intermediate attachment plate and a microscope adapter plate (Supplementary Fig. 8, described in detail below), all of which were custom designed and fabricated.

The implanted headplate (Supplementary Fig. 8) was 3D-printed using Temporary CB (Formlabs, MA, USA) and had a weight of ~100 mg. A handle, 16 mm in length, was 3D printed using black resin V4 (Formlabs, MA, USA) and attached to the headplate using UV curing dental adhesive (Charisma Flow, Kulzer GmbH, Hanau, Germany). The base of the headplate had a 5.3 mm diameter aperture below which the craniotomy for imaging was opened (details below). A hole with an independently 3D printed tapped insert at the front of the headplate was used for attaching a removable crossbar for stable fixation during surgical procedures or allowed securing a protective cover when the mice should be placed in their cage.

The intermediate attachment plate (Supplementary Fig. 8), allowed alignment of the microscope over the craniotomy and the region of interest before being rigidly attached to the headplate, again using UV curing adhesive (Charisma Flow, Kulzer GmbH, Hanau, Germany). The microscope magnetic clamp (Supplementary Fig. 8) had a central aperture that fitted tightly around the microscope, secured by a screw.

Both, intermediate attachment plate and the microscope magnetic clamp, were each equipped with 4 neodymium magnets each (ø3 mm, height 2 mm, Webcraft GmbH, Gottmadingen, Germany), providing up to 24 N holding force. Additional opposed guide fins provided stable orientation. This design allowed fast and reliable placement and detachment of the microscope during experiments. The weight of the intermediate plate, including magnets was 575 mg. The weight of the microscope magnetic clamp, including magnets was 630 mg.

All components were silanized (Rotatec bonding system, 3M ESPE, Neuss, Germany) prior to each use to improve surface adhesion.

#### Surgical procedures for fluorescent labelling of neurons with jGCaMP7f

Prior to surgery, all instruments, including the glass injection capillary, were sterilized by either autoclaving or heat sterilization. Animals were anesthetized with an intraperitoneal injection of a three component anesthetic cocktail (3K) consisting of fentanyl (50 μg/kg, Hameln pharma plus GmbH, Hameln, Germany), midazolam (5 mg/kg, Hameln pharma plus GmbH, Hameln, Germany) and medetomidine (0.5 mg/kg, Zoetis, NJ, USA). Body temperature was maintained at 37-37.5 °C with a heating pad and heater controller (FHC, ME, USA). Animal status and depth of anesthesia was monitored approx. every 15–30 min, and anesthesia maintained throughout with supplementary doses of 30–80% of the above anesthetic combination given as necessary in order to maintain withdrawal and corneal reflexes absent. The animals were then placed in a stereotaxic apparatus, the hair on the scalp removed and the skin cleaned with 70% ethanol. The right parietal bone was exposed and a burrhole drilled (approx. 500 μm diam.) at 3.0 mm posterior and 2.6 mm lateral relative to bregma. A small slit was made in the dura underlying the burrhole, and a glass injection capillary with a beveled tip containing a high titer solution of AAV1/2 coding for Cre-dependent jGCaMP7f (AAV1j7f/2.hSyn.FLEX.jGCAMP7f, Addgene, MA, USA) was advanced posteriorly into the cortex approx. 1650 μm at an angle of 25° relative to horizontal along a trajectory parallel with the saggital suture. An injection of approx. 180 nL of the virus solution was made with a nanoliter injection device (Nanoject II, Drummond Scientific Company, PA, USA). After a delay of 5 min., the tip of the capillary was withdrawn approx. 240 μm, and an injection of approx. 115 nL made. After another delay of 5 min., the tip of the capillary was again retracted approx. 240 μm, and another injection of approx. 115nL made. After a final waiting period of 5 min., the glass capillary was slowly withdrawn, the craniotomy covered with medical silicone (KwikSil, WPI, FL, USA) and the skin sutured closed using 5/0 vicryl sutures (Ethicon, NJ, USA). Animals were then administered buprenorphine (30 μg/kg, Bayer, Leverkusen, Germany) and carprofen (5 mg/kg, Zoetis, NJ, USA) for post-operative analgesia, and then a cocktail of antagonists to the anesthetic drugs (anti-3K) consisting of naloxone (11.2 mg/kg, Ratiopharm, Ulm, Germany), flumazenil (0.5 mg/kg, Hikma, Amman Jordan) and atipamezole (0.75 mg/kg, Orion Pharma, Hamburg, Germany).

#### Surgical procedures and imaging in freely moving animals

All surgical instruments and solutions used were autoclaved prior to commencement of the procedures described below. Three to five weeks after the surgery to label neurons with jGCaMP7f, animals were anaesthetized with the 3K anesthetic solution described above, and body temperature maintained at 37–37.5°C. Animal status and depth of anaesthesia monitoring procedures were as described in the section for labelling neurons with jGCaMP7f. Anesthesia was maintained with supplementary doses of 30–80% of the 3K solution. The hair on the dorsal aspect of the skull was removed and the skin cleaned with 70% ethanol. A midline incision in the skin over the parietal bones was made, the skin retracted and galea removed to expose the parietal bones, including the site of the previous burrhole. The exposed bone was then cleaned with hydrogen peroxide solution (3% by volume in sterile saline) and thoroughly washed with sterile saline. The bone was then mechanically roughened prior to application of a layer of dental adhesive (Optibond, Kerr, CA, USA). The custom made headplate (Supplementary Fig. 8, see above for details) was then fixed to the skull over the Optibond layer with dental composite (Charisma, Kulzer GmbH, Hanau, Germany). The central aperture was placed such that the burrhole from the previous surgery was located approximately centrally in the medial-lateral axis and near the anterior edge of the aperture. The skin incision was then closed firmly around the headplate using 5/0 vicryl sutures (Ethicon, NJ, USA). A circular craniotomy with a diameter of approx. 3 mm was then opened in the center of the headplate aperture, including at the anterior margin the site of the previous craniotomy. The dura was then removed and the cranial window closed using a pre-formed plug and coverslip (circular, 5 mm diam., 100 μm thickness, CS-5R-0, Warner Instruments Holliston, MA, USA,; plug custom pre-formed as a 300 or 400 μm tall cylinder of KwikSil silicone in the center of the circular coverslip after the description in^23^).

The animal was then transferred to the miniature microscope, which was mounted on a navigation stage for locating an appropriate position for imaging within the cranial window. The navigation stage consisted of a micromanipulator (MP-285, Sutter Instruments, CA, USA) to which the microscope was mounted using a custom made mount. The mount included two angular kinetic mounts (GN05/M and GN1/M, Thor Labs, NJ, USA) allowing adjustment of tilt with respect to the coverslip and cortical surface. Once a target population of neurons had been located, the intermediate attachment plate (already mounted to the miniature microscope) was attached to the headplate with dental composite (Charisma Flow, Kulzer GmbH, Hanau, Germany), the microscope removed and the animal administered the anti-3K cocktail and analgesic combination as described above.

Freely moving experiments were subsequently conducted from 2 to 10 days after opening the cranial window and positioning the microscope. At the commencement of each recording session, the animal was taken from its home-cage, the head gently restrained by holding the handle on the back of the headplate, and the microscope placed onto the intermediate attachment plate. Placement of the microscope was rapid, taking in the order of a few seconds. After microscope placement, the animal was placed onto the linear track and allowed to explore. Candidate populations of neurons were located by navigating with the remote z-drive mechanism, and then data acquired. Frame rate and acquisition parameters were 273*280 pixels per frame, 10.6 frames per second. At the end of a recording session, the animal was again gently removed from the linear track, the screw on the microscope mounting plate unfastened and the microscope removed. This mechanism allowed the microscope, and particularly the objective which projected through the microscope mounting plate and intermediate attachment plate and made direct separation of the plates difficult, to be lifted vertically away from the headplate without having to apply the force required to separate the magnets. With the microscope removed, the magnets holding the microscope mounting plate and intermediate attachment plate could be separated with the animals head restrained and supported again using the tab on the headplate, and the intermediate attachment plate removed. The central aperture of the intermediate attachment plate was then closed using a cap that was also fitted with matching magnets, after which the animal returned to its home-cage.

#### Linear track and animal position and orientation tracking

The linear track was 96.0×9.7 cm^2^, surrounded by a rim 2 cm in height and raised approx. 110 cm off the floor. The animal was free to explore and move around the track during experiments. Animal position and head orientation were tracked as described in ^24^. In brief, 3 struts were mounted onto the body of the miniature microscope, with 3 infrared (IR) LEDs (940 nm, SFH 4053, Osram) mounted on each. Images of the IR-LEDs were acquired at 200 Hz by 4 calibrated and synchronized digital cameras (acA1300-200um, Basler AG, Ahrensburg, Germany) mounted above the track. The cameras were equipped with 8.5 mm/F1.5 objective lenses (Pentax, Ricoh Imaging, Tokyo, Japan) to facilitate coverage of the track and IR bandpass filters (LP 780-40.5, Midwest optical systems, IL, USA) to facilitate automated tracking of the LEDs in the images. The exposure active signals from the overhead tracking cameras as well as the frame synchronization signal from the miniature microscope were fed into an analog-digital converter (Power 1401, Cambridge electronic design, Cambridge, UK) and recorded with Spike-2 software (Cambridge electronic design, Cambridge, UK) for synchronization of behavioral tracking and multiphoton imaging data. Determination of animal position and head orientation were performed using the method described in detail in^24^, initialized by a pre-detection step using multitrackpy (https://github.com/bbo-lab/multitrackpy). Mounted directly next to each of the above cameras was a second camera (acA1300-200um, Basler AG, Ahrensburg, Germany), equipped with an 8.5mm/F1.3 lens (Edmund optics, New Jersey, USA) for simultaneously recording the animals position on the track at the visible light level. All cameras were calibrated using the calibration procedure described in^7^.

#### Histology

At the termination of the imaging experiments animals were deeply anesthetized with ketamine (100 μg/kg) and medetomidine (200 μg/kg), and perfused transcardially with 0.1 M phosphate buffer (PB) followed by 4%formaldehyde solution (Roti-Histofix, Carl Roth, Karlsruhe, Germany). The brain was then removed, post-fixed at 4°C overnight in the same formaldehyde solution and then transferred to 0.1 M PB. Sections of 100 μm thickness were then cut on a vibrating microtome (Leica VT1000S, Wetzlar, Germany) and mounted in fluoroshield (Sigma, MO, USA). Images were acquired on an inverted microscope (Nikon Ts2R-FL, Tokyo, Japan).

### Microscope and optical setup

#### Excitation laser setup

A non-collinear optical parametric amplifier (NOPA), pumped with an Ytterbium fiber laser (Spirit-NOPA, Spectra Physics, CA, USA) was used to produce laser pulses at 1300 nm center wavelength with a maximum average power of 3 W (3 μJ at 1 MHz). Compressed pulses (built-in compressor, 3 mm of ZnSe) were sub-50 fs at the output of the laser. We used a half-wave plate (AHWP05M-1600, Thorlabs, NJ, USA) mounted on a stepper-motor controlled rotator (G065118000, Qioptiq, MA, USA) with a polarization beam splitter (PBS104, Thorlabs, NJ, USA) to control beam intensity. To compensate dispersion introduced by the fiber we used a two-prism sequence with additional bulk silicon described previously^6^. In brief, to compensate third order dispersion [TOD], a double-path two-prism sequence compressor with Brewster-angle silicon prisms at 32° apex angle (4155T724, Korth Kristalle GMBH, Altenholz, Germany) were used. We used a custom-designed second prism with a with larger base of 56 mm to increase the compensation range. Additional anomalous group-velocity dispersion (GVD] generated by the prism-sequence was compensated using additional bulk silicon. To pre-compensate the dispersion of the 1.5 m of fiber used in the experiments in the current study we used an inter-prism distance of 35 cm and an additional 10 cm of silicon. The beam was coupled to the previously designed custom hollow-core fiber^6^ with a 10 mm focal length achromatic lens (AC050-010-C-ML, Thorlabs, NJ, USA). A half-wave plate (**AHWP05M-1600**, Thorlabs, NJ, USA) was used to control the linear polarization orientation of the laser with respect to the fiber structure and a detuned 1:1 relay telescope enabled optimization of the coupling efficiency of the fiber.

#### Miniature optics design and optimization

We used OpticStudio 14.2 (Zemax Europe, Bedford, UK) to design the full optical system for the three-photon miniature microscope. The system was composed of a collimation lens, scanning optics (scan lens system and tube lens) and an imaging objective lens (Supplementary Fig. 9). The objective lens was designed first, and then all the other parts were optimized as part of the full system with objective lens parameters kept constant. The primary design parameters for the objective lens were: large working distance (> 1.5 mm), high collection NA (> 0.9), excitation NA up to 0.6 and large field of view (> 400 μm, at 0.4 excitation NA). As three-photon excitation requires short pulses, chromatic aberrations constitute a particular concern for microscopes employing it. To account for it, three wavelengths were used with equal weight for the simulations (1250 nm, 1300 nm, 1350 nm). The polychromatic Strehl ratio (SR) was used as the main quantification of the optical performance.

Compensating field curvature requires complicated, large and heavy objective lenses^25^. A fully uncompensated field curvature led to about 10 μm axial distance between the axial focal point and the outer part of the FOV. This parameter was left floating and uncompensated to keep the objective lens as small as possible, and because such a field curvature was not deemed to be an obstacle to measure neuronal activity in the intact brain, as 10 μm distributed over 200 μm of half-field-of-view produces a small z-slope in relation to a z-resolution > 10 μm.

To keep the weight and the size of the system minimal, several approaches were used during the design process. First, the effective focal length (EFL) of the objective was decreased compared to the previous design (1.8 mm from 3 mm). This resulted in much smaller beam diameters employed to achieve the same NA (3.6 mm compared to 6 mm for an NA of 0.9) and allowed the use of much smaller and thinner lenses. The focal lengths of the scan and tube lenses were also minimized to shorten the total beam path. The limitation to this approach is the proportionally larger scanning angles (compared to beam diameter decrease) required to scan the same field of view (FOV). Focal lengths were systematically explored and the best balance (achieving minimal weight and size) was selected for the design.

The same guidelines were used to design scanning optics and the collimation lens. A 2.0 mm diameter MEMS scanner (A7M20.1-2000AU-LCC20-C2TP, Mirrorcle, CA, USA) was selected, featuring +/- 4.5 mechanical degrees of scanning range. To minimize the manufacturing costs the tube and scan lenses were constrained to contain the same achromat. The tube lens is the achromat itself and the scan lens is composed of the achromat and a plano-convex (PCX) lens of either 15, 12 or 9 mm EFL (3 mm diameter, EFL of 15 mm, 12 mm and 9 mm, PCX lenses, NIR II, Edmund Optics, NJ, USA) to provide a range of magnifications. The focal length of the achromat was chosen to be 6.68 mm, to provide 0.4 excitation NA with the scan lens containing a PCX of 15 mm EFL. In the case of 12 or 9 mm EFL PCX lens the excitation NA was 0.43 and 0.48 respectively.

Additional constraints were provided by the requirement for imaging under defocus conditions to implement remote focusing (Supplementary Fig. 9). To minimize the system complexity and size, an unfolded beam path was chosen with the defocus provided by a varying distance between the hollow-core fiber (HCF) and the collimation lens. The distance from the collimation lens to the MEMS scanner was the EFL of the collimation lens to provide constant excitation NA over the Z range. The combination of the collimation lens EFL and the NA of the HCF with the magnification of the scan-tube lens system give the final excitation NA. The design parameters of the Z-drive were: a Z-range of >600 μm (covering a substantial part of the mouse cortex in depth) with >0.4 excitation NA and >400 μm FOV.

Achieving an NA of 0.4 exacts a beam diameter of 1.6 mm at the back focal plane (BFP) of the objective. A FOV of ±200 μm requires a scan angle of arctan(±0.2/1.8) = ±6.34° at the BFP. Considering the deflection properties of the MEMS scanner a magnification of 2o4.5°/6.34 ° = 1.42 is required in the scan-tube lens system. With a magnification of 1.42, to achieve a beam diameter of 1.6 mm at the BFP of the objective, a 1.6 mm/1.42 = 1.13 mm beam diameter is needed at the MEMS, which, provided an estimate of 0.09 NA of the HCF, requires a collimation lens of 6.29 mm EFL. We used a combination of two EFL 12.0, PCX lenses (#67-444, Edmund Optics, NJ, USA) to decrease aberrations due to defocus in the collimation lens. The achromat used as tube and scan lens in combination with a PCX lens with an EFL of 15 mm yielded a magnification of 1.42 of the scan-tube lens system (EFL of 6.3 mm). All previous values were kept constant during the optimization of the design.

A dichroic mirror (T875spxrxt, AHF analysentechnik AG, Tübingen, Germany) was used to separate the excitation beam path and the emitted fluorescence. A condenser lens with an EFL of 5 mm was used (Throl GmbH, Wetzlar, Germany). To produce a high performance miniature infra-red filter, we used an existing filter with an optical density (OD) of 7 in the range 900–1600 nm (T875spxrxt-1800-UF1, Chroma Technology Corp, VT, USA), which was cut in circular pieces 4 mm in diameter. The resulting pieces were separated in two groups and ground down to a thickness of 0.3 mm starting from either side of the filter. Pairs were cemented together (one piece from each group) to obtain a 4 mm diameter filter with the original performance and thickness of 0.6 mm. A dichroic mirror (DMLP490R, Thorlabs, NJ, USA) separated detected light into two channels. This mirror was cut into rectangular pieces of 2 mm × 3 mm and ground down to 0.3 mm thickness. We used square, 1.3 mm side, silicon photo-multipliers (S13360-1375PE, Hamamatsu, Hamamatsu City, Japan) as detectors on-board of the microscope (see below).The resulting system is shown in Supplementary Figure 9.

For the experiments with the electrically tunable lens (ETL,) we used an electrowetting lens (A-25H1-D0, Corning, NY, USA), with the compatible driver (USB-M Flexiboard, Corning, NY, USA) and a relay lens system composed of 4 lenses, 2 with EFL 12 mm and 2 with EFL 9 mm ((#67-444 and #67-443, Edmund Optics, NJ, USA) as shown in Supplementary Figure 3.

#### Microscope and Z-Drive manufacturing

The structure of the microscope body, implants and lenses mounts, including objective, were designed with Inventor (Autodesk GmbH, München, Germany). A 3D printer (Form3, Formlabs) was used to produce the microscope body in black resin V4 (Formlabs, MA, USA) and the headplate in a biocompatible, glass-filled resin (Temporary CB, Formlabs, MA, USA). All the optics and the objective frame, made of titanium, were supplied by Throl GmbH (Throl GmbH, Wetzlar, Germany).

The HCF was passed through a jacket (FT900Y, Thorlabs, NJ, USA). The end of the jacket from the laser to fiber coupling side was glued in a plate for a 30 mm cage system (Thorlabs, NJ, USA). A syringe was then glued on this plate, centered and aligned along the axis of the cage. To prevent the fiber from buckling during sliding the bare fiber was passed through that syringe first, then through a smaller syringe, with an outer diameter that matched the inner diameter of the first syringe, such that the second syringe could slide inside the first (Supplementary Fig.11). The bare fiber was glued inside the second syringe, so displacement of this syringe inside the cage system was displacing the bare fiber inside its jacket. The jacket was rigidly attached to the microscope body, so the displacement of the fiber inside the jacket resulted in variation of the distance of the fiber tip to the collimation lens from < 1 mm to > 14 mm. The actuation was performed using a stepper motor (ZST213, Thorlabs, NJ, USA). On the microscope side, the fiber tip was glued inside a ceramic ferrule of 2.5 mm diameter (CF128-10, Thorlabs, NJ, USA) which slid inside a corresponding structure on the microscope (supplementary fig. 9), that was 3D-printed with the diameter adjusted with a reamer to minimize sliding friction.

#### Fluorescence detection, control electronics and software

Scanimage 5 (Vidrio Technologies, VA, USA) with custom FPGA code featuring the triggered acquisition was used to control the microscope. The MEMS scanner was operated in the resonance mode in the fast axis at 1.5 kHz period providing 3000 lines per second using bi-directional scanning.

This, combined with a 1 MHz excitation laser pulse rate and 82% of imaging fill fraction, resulted in a resolution of about 273 pixels * 280 lines at 10.6 Hz. The field of view during the activity data collection was a square of up to 300×300 μm^2^. The driving signals from Scanimage software were sent to the MEMS driver (BDQ_PicoAmp_4.6, Mirrorcle, CA, USA).

#### Environment light

The environment of the track was homogeneously illuminated using 6 125 cm long 24V RGBW LED strips with 910 lm/m and 8 125 cm long 12V white LED strips with 700 lm/m (both LED strips from PowerLED, Berkshire, United Kingdom), arranged equidistantly in a patch of 125×60 cm^2^ at a distance of 150 cm above the track.

The strips where switched on and off in synchrony with the line signal of the microscope using a custom circuit based on TIP120 Darlington transistors.

### Data analysis

#### Alignment of single frames for motion correction

Motion correction was performed using a custom MATLAB port of the image registration component of suite2p^26^ (parameters: align_by_chan=1, nimg_init=200, batch_size=5000, smooth_sigma=1.15, maxregshift=0.1, nonridid=0), involving a displacement estimation using phase-correlation and subsequent rigid frame shift. In addition to the corrected movie, the algorithm yields the per-frame displacements in pixels for horizontal and vertical direction respectively, which have been used for the quantification of brain movement.

#### Extraction of fluorescence signal

The fluorescence *F_t_* in the *t*^th^ frame for a given neuron was extracted by calculating the per-frame average of pixel values in a region of interest (ROI). ROIs where manually drawn around the somata of cells in focus and linked over a continuous recording session of multiple files.

#### Calculation of ΔF/F_0_

The baseline fluorescence *F*_0,*t*_ was defined as the mean of the lowest 20% of fluorescence values within a window of maximally 107.63s around the *t*^th^ frame (if not truncated by beginning or end of the file) after applying a Gaussian filter with a standard deviation of 282ms, and *ΔF_t_/F*_0,*t*_ = (*F_t_* – *F*_0, *t*_)/*F*_0,*t*_.

#### Spike detection

Spike detection was performed using MLspike^21^ (available at https://github.com/MLspike/spikes,%20commit%2032fb84e) on dF/F_0_, using parameters F0 = [-0.5 1.3], a = 0.034, pnonlin = [0.85 −0.006], drift.parameter = 0.015, finetune.sigma = 0.08, algo.nc = 50, algo.nb = 50. Since no parameters for jGCaMP7s are available, these parameters were based on the ones for GCaMP6f and manually inspected for consistency with the fluorescence signal. While consistency of active an inactive period is well maintained between transients and inferred spikes, absolute numbers of spiking rates are to be interpreted with care.

#### Shuffling of spikes

To gauge the significance of the observed light-dark responses, we artificially shuffled the recorded frames to create a control group of random firing. To maintain the original structure in the firing, we shuffled as follows: First, we created a list of inter-spike intervals (ISI), consisting of the intervals between spike times, and the sum of the beginning and end intervals between recording start and end and the first and last inferred spike, respectively. Subsequently, we separated this list into a list of short (<= 3*dt) ISIs and list of the remaining longer ISIs. We then created a list of spike train lengths, that is, numbers of spikes that are only separated by short ISIs. The list of long ISIs was summed and the total time was randomly redistributed to the same amount of ISIs.

Finally, we constructed a new, randomized spiking pattern by randomly sampling (without replacement) from the lists of train lengths, short ISIs (for the ISIs within a spike train) and long ISIs (for the ISIs between spike trains). The initial interval before the first spike train was drawn from the list of long ISIs and subsequently multiplied by a random number between 0 and 1. We used this method 100 times on the original data to create 100*n randomly firing neuron.

## Acknowledgments

We thank Ruth Pohle for assistance with editing. We thank U. Czubayko and Jeanine Klesing for expert technical help. Funding from Stiftung caesar and Max Planck Society.

## Author contributions

Experimental design AK, DJW, YG, JNDK, microscope hardware and software development AK, JS. Animal preparation DJW, YG. Data collection AK, DJW, JS, KMV and JNDK. Analysis design and implementation AK, DJW, KMV and JNDK. Manuscript preparation AK, DJW, JS and JNDK.

## Data availability

On publication, the data will be available via a Dryad data repository. Zemax files for the optics developed will be available on publication.

## Code availability

No custom code central to the results presented in the manuscript were developed.

---

**Supplementary Figure 1.**
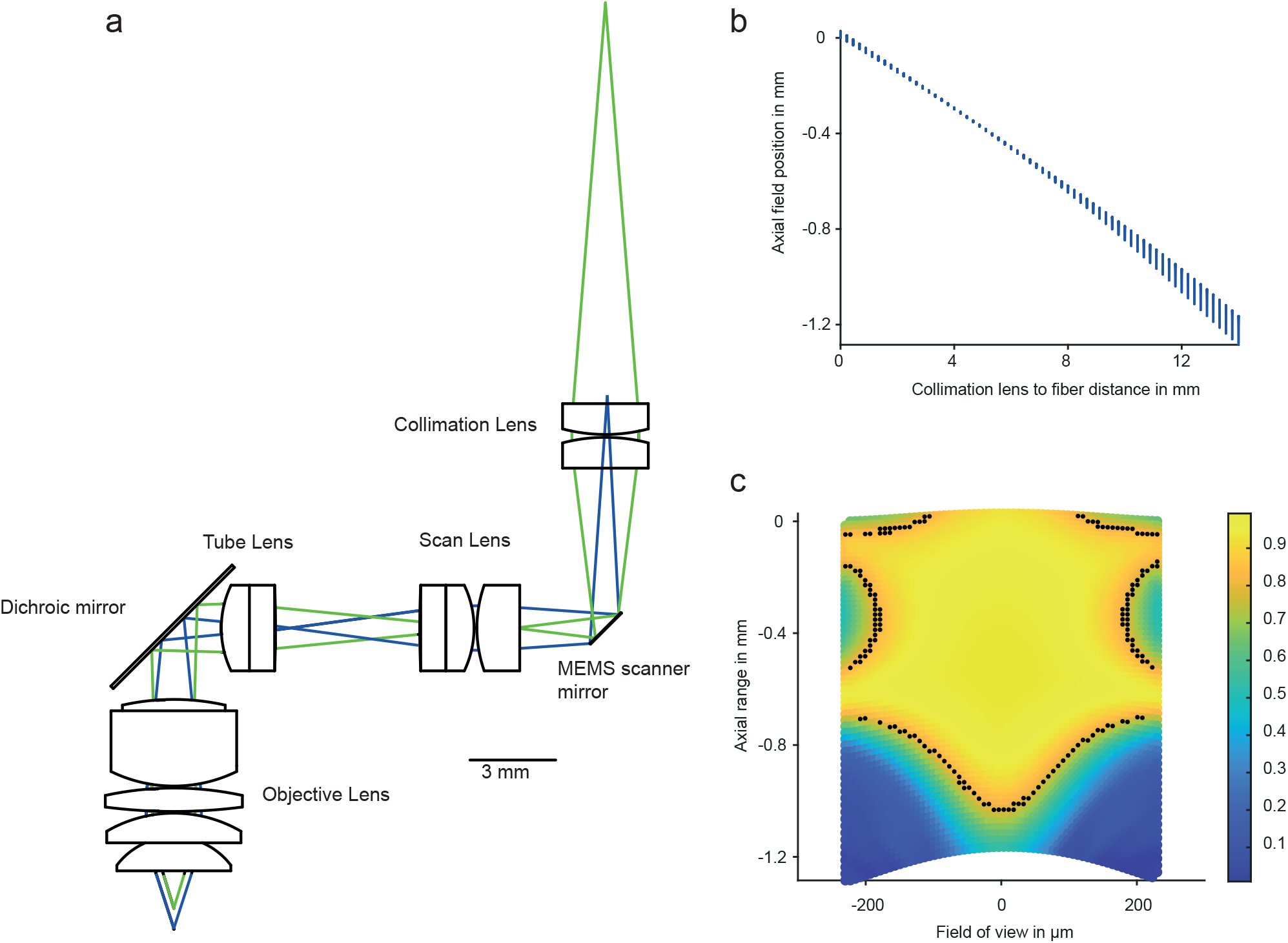
Optical design and performance simulation in Zemax. **a**, optical layout of the microscope. Green and blue rays illustrate extreme position of the fiber tip leading to minimal and maximal working distance respectively. **b**, imaging depth relative to the maximum working distance as function of the fiber tip to collimation lens distance from Zemax simulation. Note that for each collimation lens to fiber tip distance there is a range of imaging depths because the objective is not corrected for flat field. **c**, simulated optical performance measured by polychromatic Strehl Ratio (SR, color-coded) over the field-of-view and the Z-range achievable by changing the fiber-collimation lens distance. Black dots delimit the diffraction limited imaging area (SR > 0.8).

**Supplementary Figure 2.**
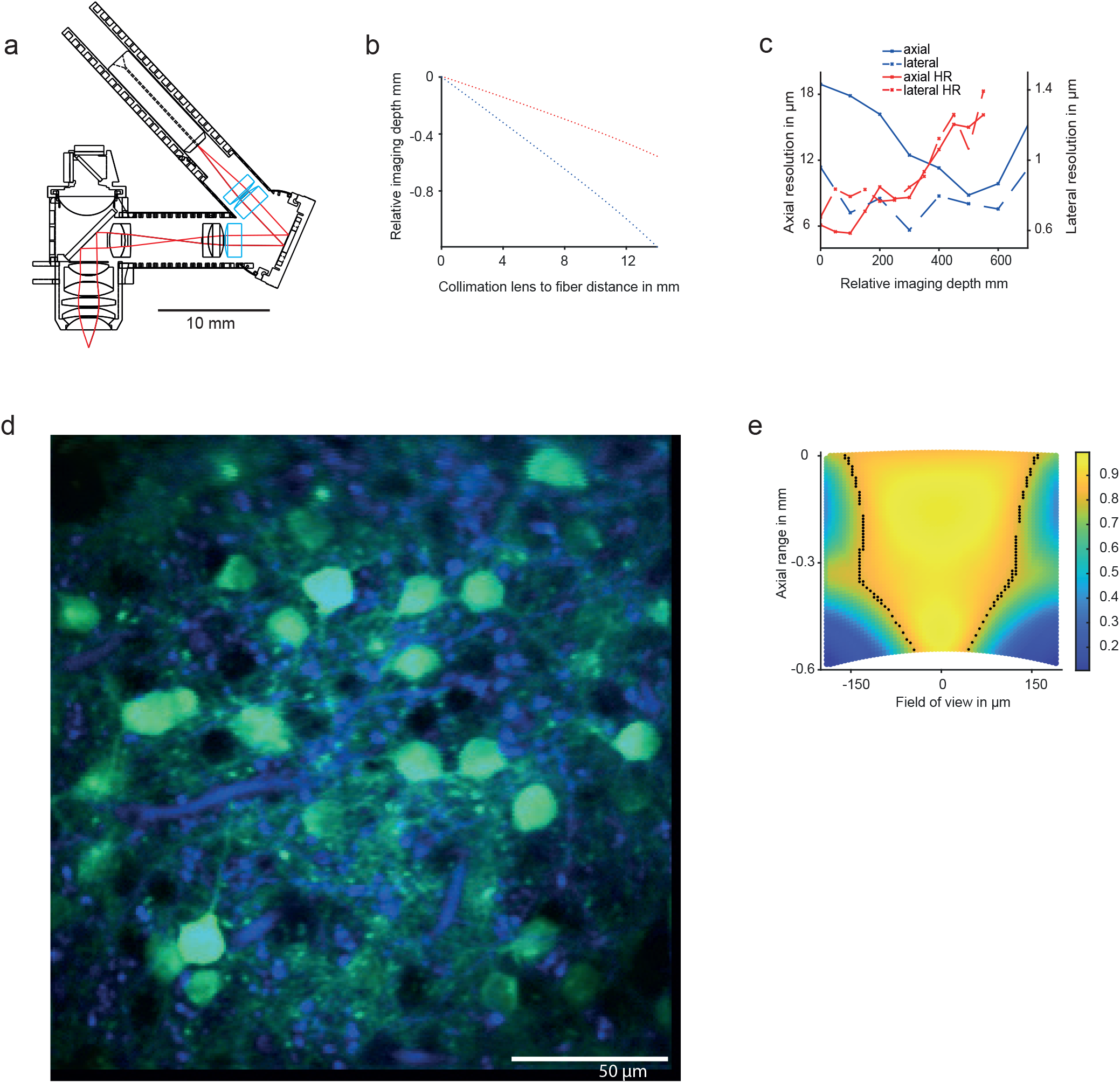
The choice of commercial lenses determines the interplay between resolution, field-of-view and Z-range. **a**, optical layout of the microscope showing the commercial of-the-shelf (COTS) lenses (blue). COTS lens selection determines the microscopes resolution, field of view and z-range. **b**, simulation of the z-range for the high z-range microscope presented in the main text (blue) and a high resolution version with different set of COTS lenses (red). **c**, measured axial and lateral resolutions of the high z-range and high resolution versions of the microscope. **d**, example field-of-view of the high resolution version microscope in layer 4 of an anaesthetized mouse. **e**, simulated optical performance for the high resolution version measured by polychromatic Strehl Ratio (SR, color-coded) over the field-of-view and the z-range achievable by changing the fiber-collimation lens distance. Black dots delimit the diffraction limited imaging area (SR > 0.8).

**Supplementary Figure 3.**
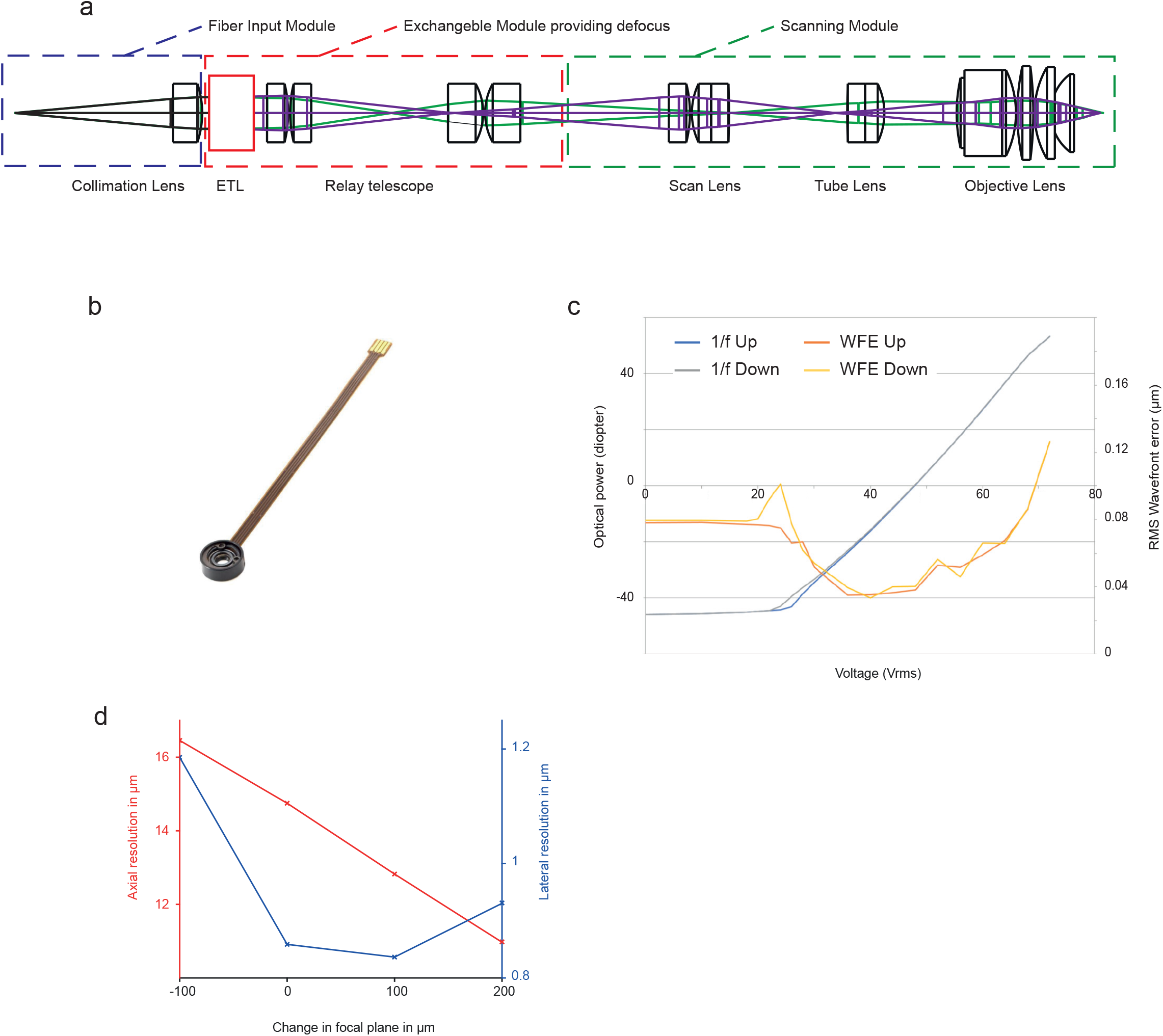
Optical design of the microscope employing an electrically tunable lens for defocus. **a**, linear optical layout of the microscope using an electrically tunable lens. Violet and green rays show the beam in two extreme defocus positions giving minimal and maximal working distance respectively. **b**, photograph of a high dynamic range electro wetting electrically tunable lens. c, optical power and RMS wavefront error of the electrically tunable lens as function of voltage. **b** and **c**, reproduced from Varioptic lens documentation with permission from Corning. **d**, measured axial (red) and lateral (blue) resolutions of the electrically tunable lens based version of the microscope.

**Supplementary Figure 4.**
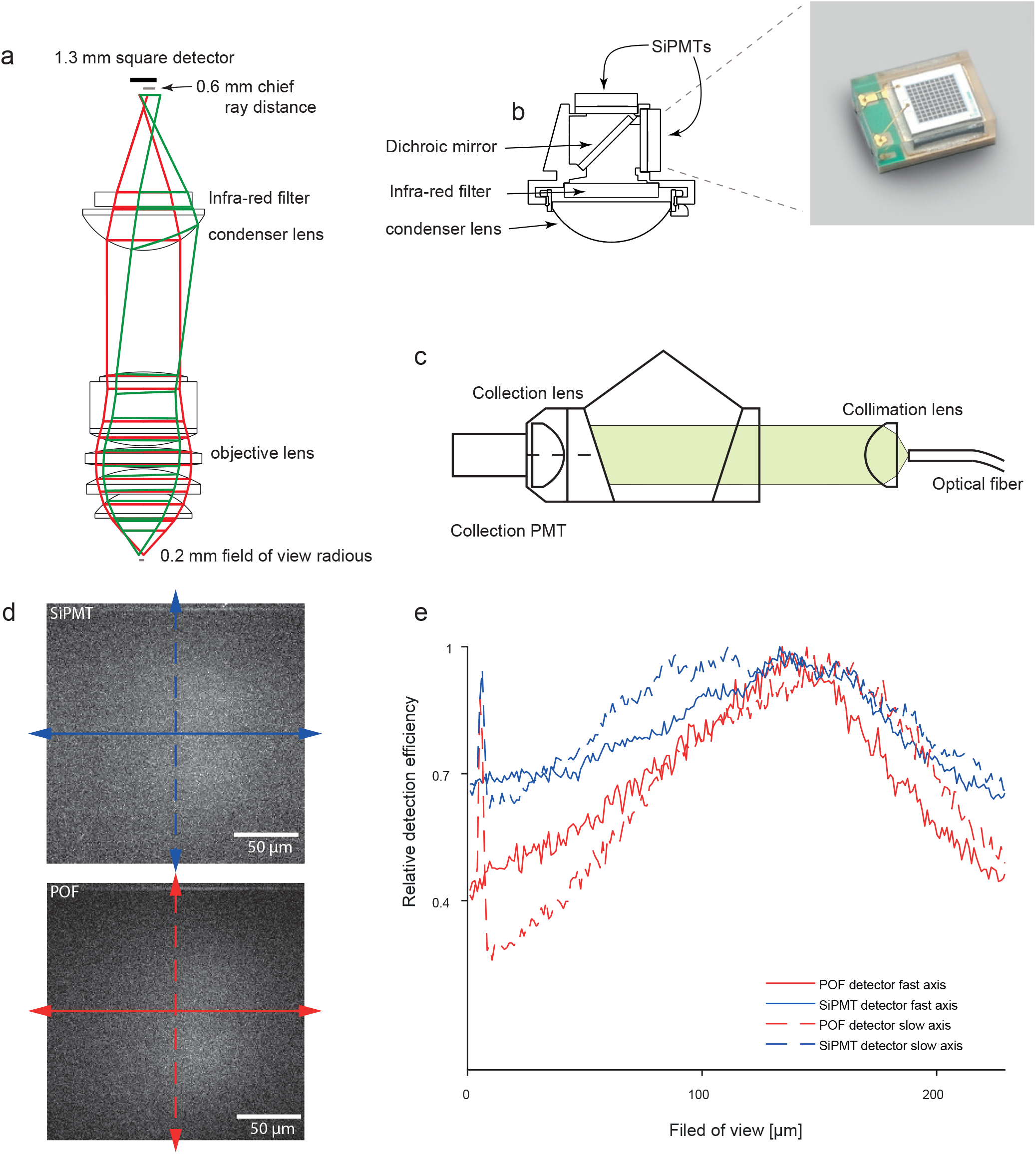
Detection system utilizing SiPMTs. **a**, optical simulation of the detected fluorescence with 0.9 collection NA and 200 μm radius field-of-view. Gray lines represent one radius of field-of-view (bottom) and distance between chief rays of fluorescence detected on-axis and 200 μm off-axis at the detector plane (top). The detector surface is also shown in black, for comparison. Note that some vignetting appears in the extreme region of the field-of-view. **b**, mechanical layout of the 2 channel detector system (left) and photograph of a SiPMT detector (right). **c**, detector system layout employed for detecting fluorescence through the plastic optical fiber. **d**, average of 200 frames of imaging fluorescein solution under identical conditions with the SiPMT based detector system (top) or plastic optical fiber baser detection system (bottom). Both images are on the same look-up table. The axes shown are used to calculate projection profiles in e. The bright stripe in the top region of both images is the flyback period. **e**, relative detection efficiency through the axes shown in d, for the SiPMT- (blue) and POF-based (red) detection. Each curve has been normalized to its maximum. The maximum detection efficiency for both schemes is compared in Figure 2b. Note the peak on the side of the field of view is due to the flyback.

**Supplementary Figure 5.**
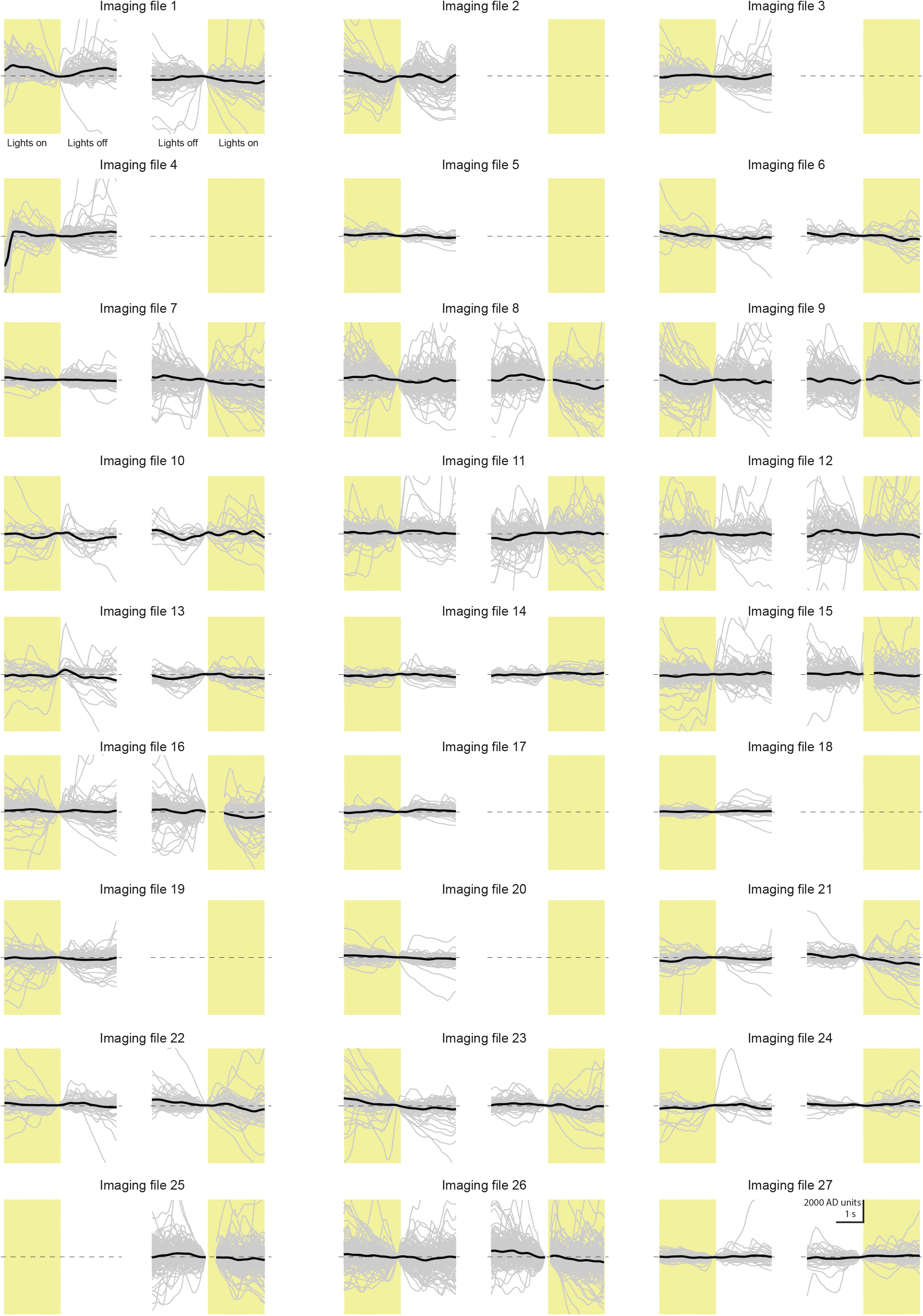
Turning ON or OFF the environment light does not produce systematic offset in the fluorescence recording. Compiled raw jGCaMP7f fluorescence traces from all neurons in all the data files over environmental lighting transitions from on to off (left side of each panel) or off to on (right side). Gray traces are traces from individual neurons, black traces represent the average. Yellow boxes represent periods with the environmental lighting on, white background represents periods with the environmental lighting off. Scale bar in the bottom right panel applies to all panels.

**Supplementary Figure 6.**
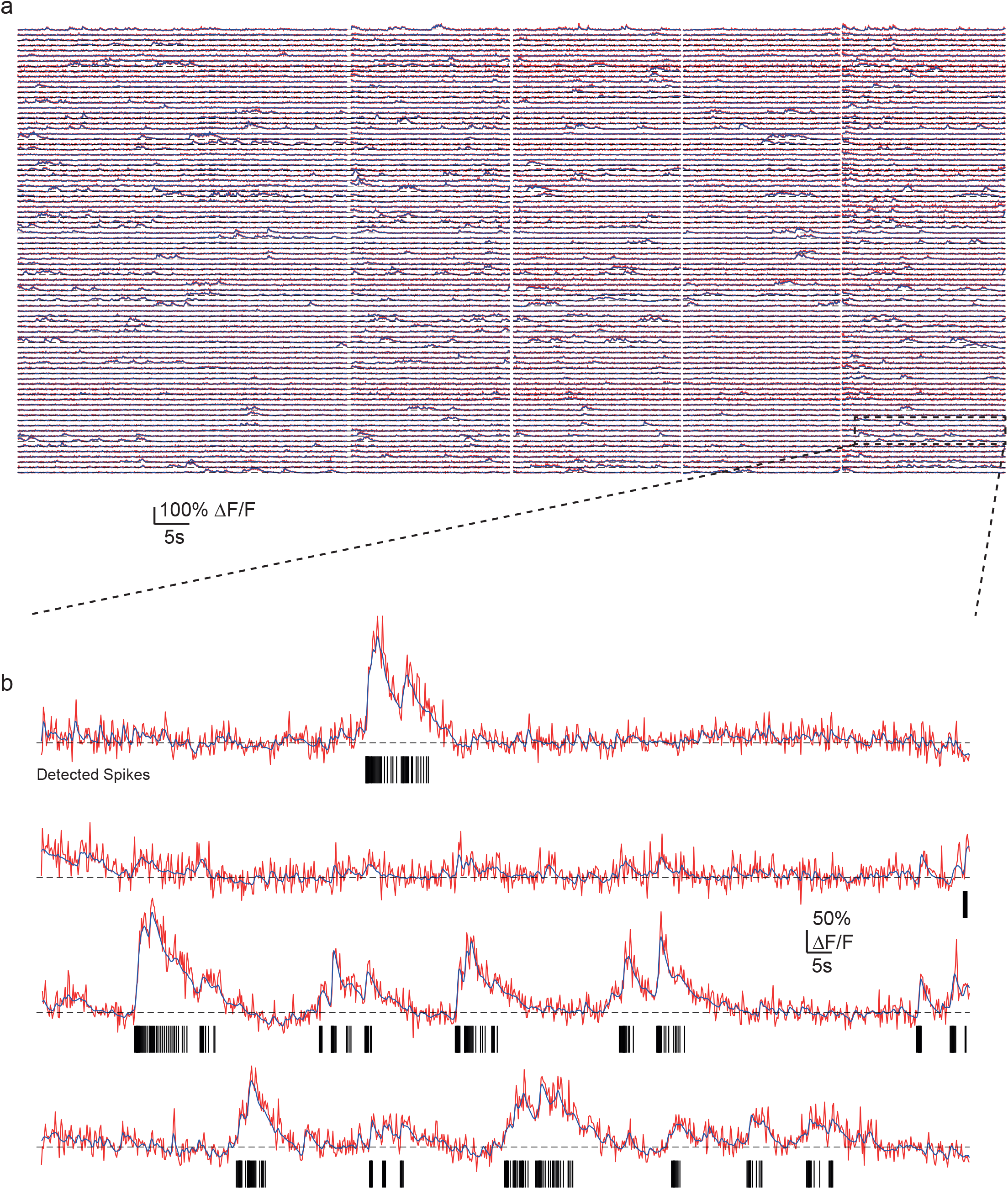
Example performance of de-noising and spike finding algorithms on acquired data. **a**, neuronal activity from all neurons during one recording session in cortical layer 6, showing raw traces from each neuron (red), estimated baseline for each neuron (black dashed) and the de-noised output of DeepCAD (blue)(Li, Zhang et al. 2021). **b**, enlarged portion of indicated region in a. Each black Vertical lines represent inferred spike times from ML spike (REF). Color scheme as in a.

**Supplementary Figure 7.**
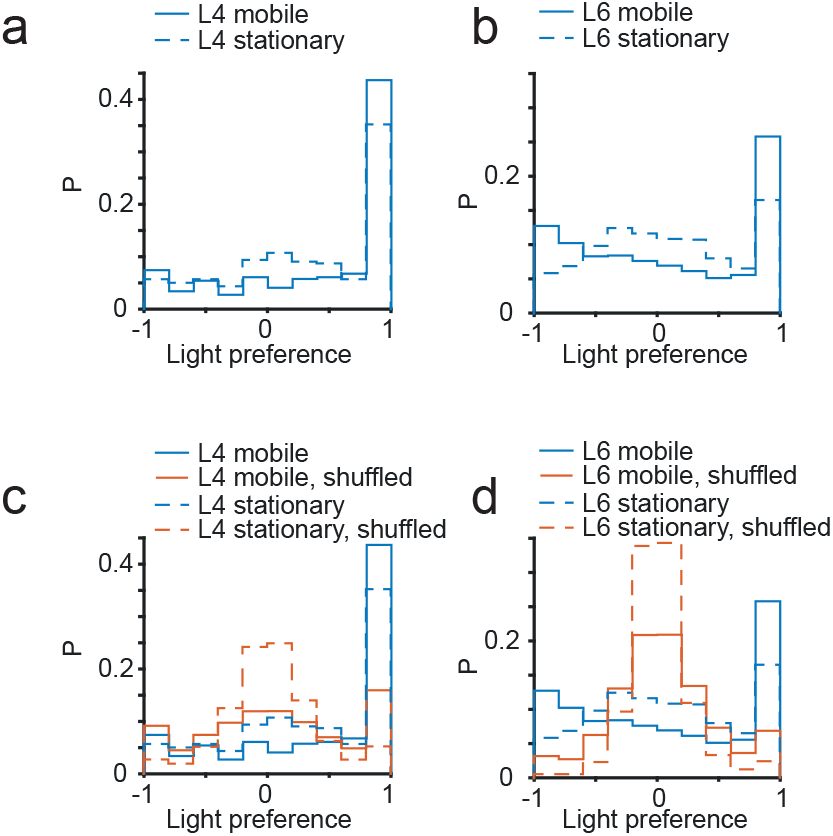
Light preference of neurons when the animal is mobile or stationary. **a**, distribution of light preference indices for L4 neurons from all datasets (N=298 neurons from 9 datasets from 3 animals) with animal mobile (solid) or stationary (dashed). **b**, as for a, but for all neurons from all datasets in L6 (N=880 neurons from 9 datasets from 4 animals). **c**, same as a, with superimposed shuffled data (see Methods). **d**, same as b, with superimposed shuffled data.

**Supplementary Figure 8.**
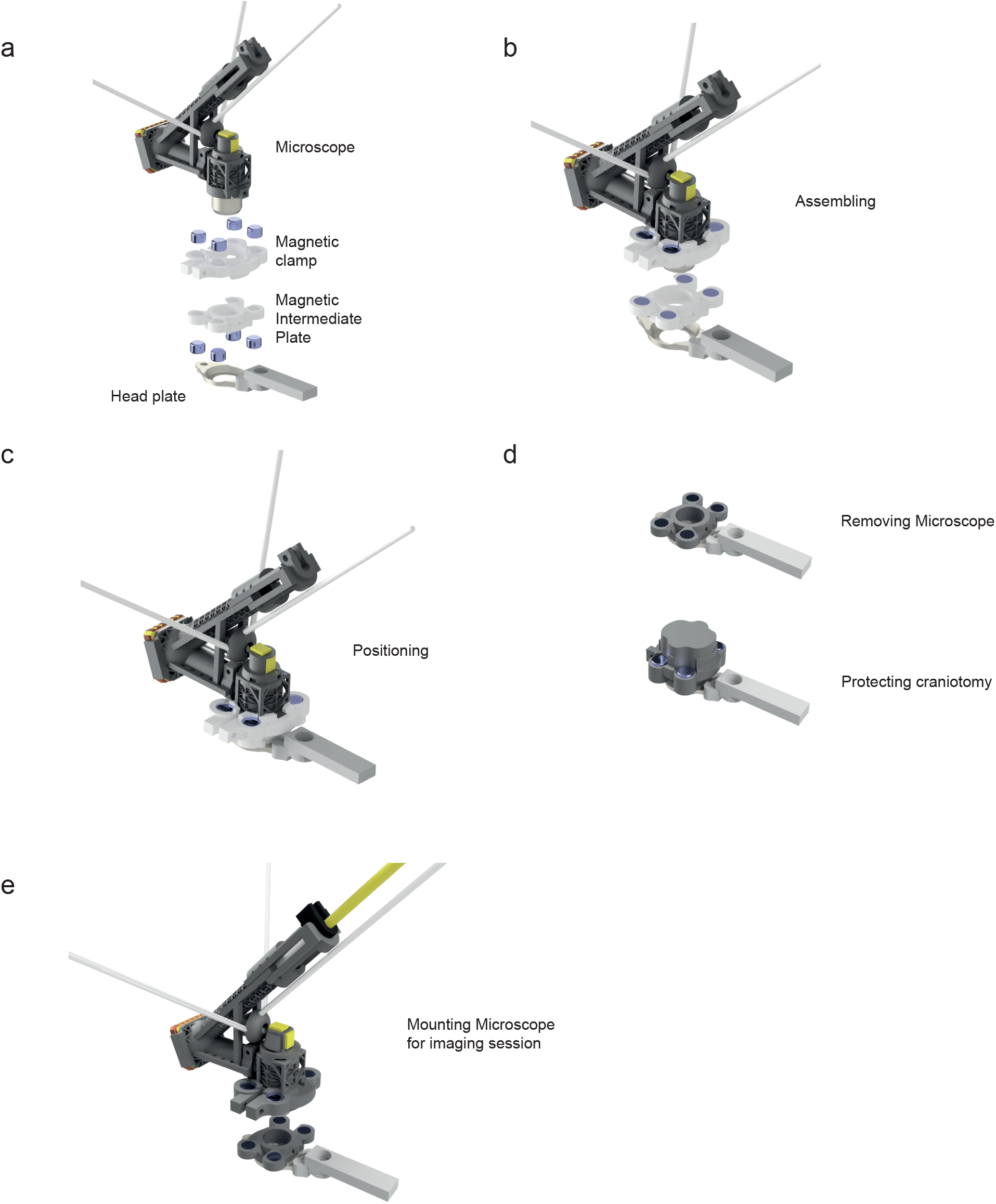
Procedure for mounting the microscope on the headplate. **a**, exploded view of the mounting parts. **b**, microscope mounted in the magnetic clamp prior to attachment of the magnetic intermediate plate. **c**, microscope with magnetic clamp and magnetic intermediate plate attached as mounted into the 3D positioning system. The positioning system allows navigation around the cranial window and positioning over a clear field-of-view of neurons. **d**, after positioning, the intermediate plate is fixed to the head plate, the microscope removed and the intermediate plate covered with a protective cover. **e**, microscope and magnetic clamp assembled for mounting the microscope for an imaging session. See also Supplementary movie 5.

**Supplementary Figure 9.**
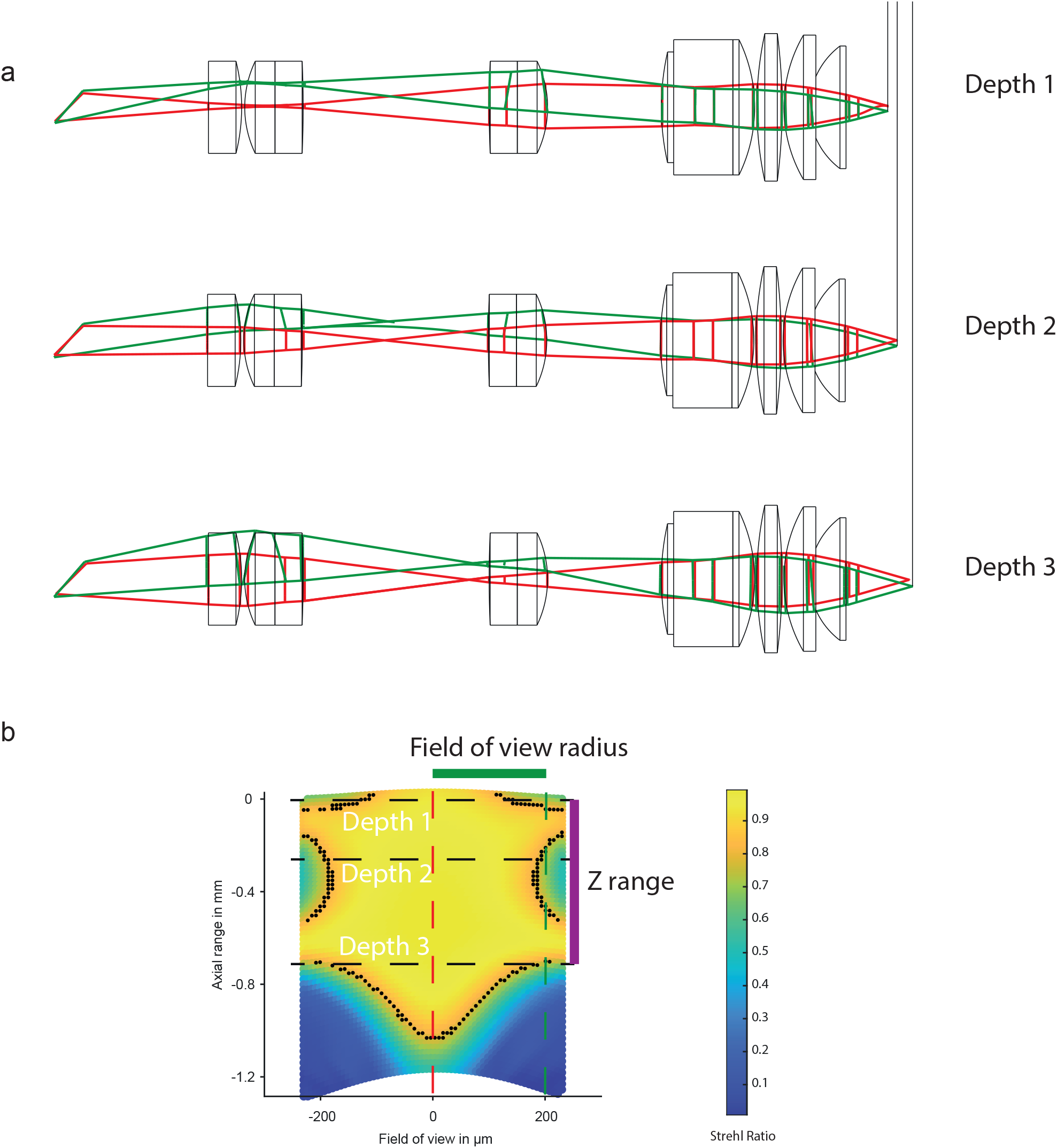
Initial step in the optical design: scanning module under multiple defocus conditions. **a**, optical layouts illustrating three configurations with different defocus applied to the input beam. The first surface modeled (left of image) is the MEMS scanner mirror. On-axis and maximal mirror deflection (extreme side of imaged area) beams are shown in red and green respectively. This system corresponds to the original optimization conditions. **b**, simulated optical performance measured by polychromatic Strehl Ratio (SR, color-coded) over the field-of-view and the z-range achievable by changing the fiber-collimation lens distance. Black dots delimit the diffraction limited imaging area (SR > 0.8). The on-axis and maximum deflection for field-of-view positions are indicated by the red and green dashed lines, and the three depths by the black dashed lines.

**Supplementary Figure 10.**
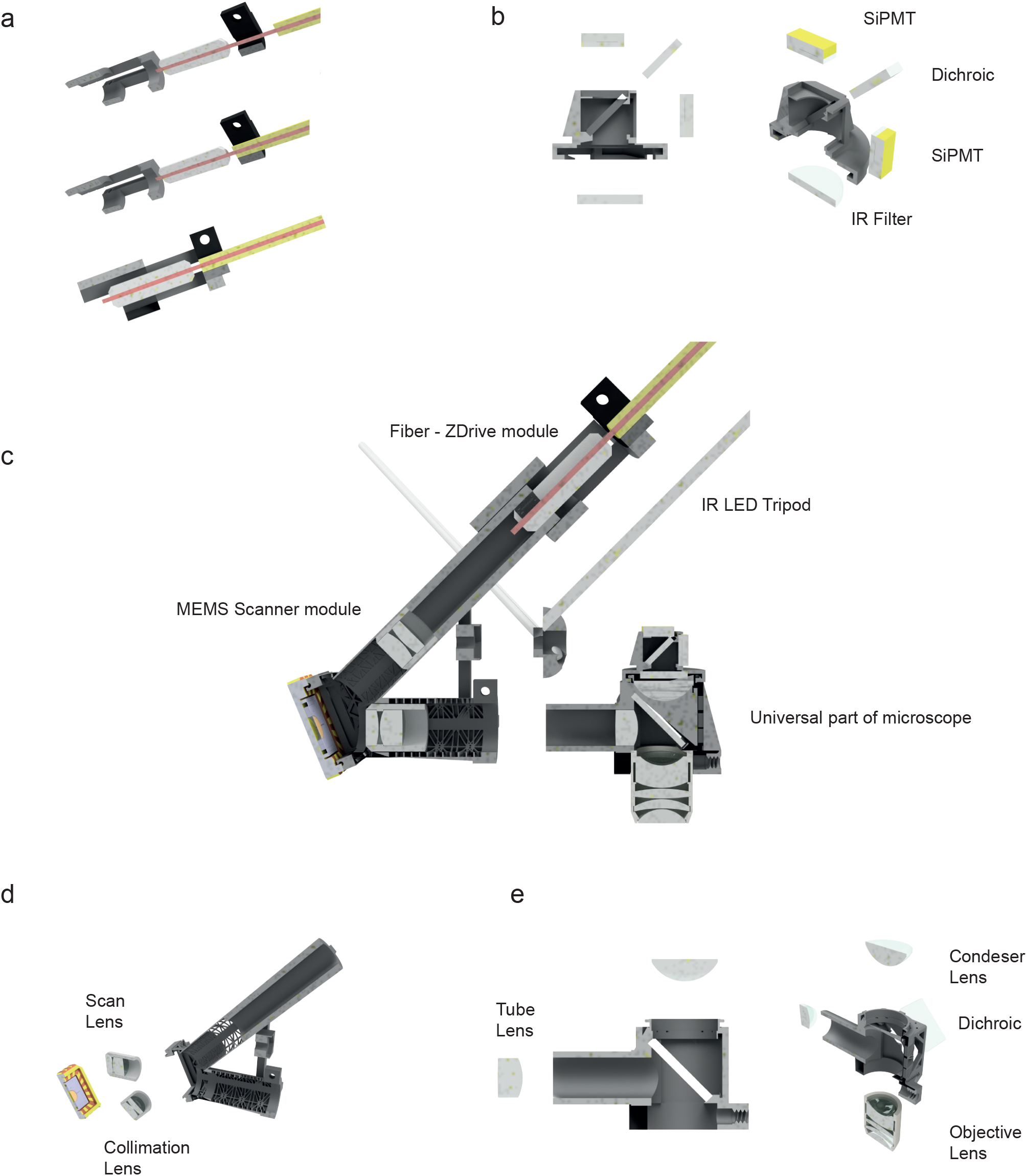
Building the microscope: profile and oblique views on sections of all microscope components. **a**, fiber z-drive module mounting: first the ferrule is inserted, then the jacket is pressure fitted in the attachment part, finally the part is attached together. **b**, detector system. First the dichroic mirror is inserted through the top surface. Optional fluorescence filters are glued on top of the detectors, and then the detectors are glued inside their compartments. Finally the infra-red (IR) filter is inserted into the bottom aperture. **c**, the final microscope assembly step is to join together the MEMS scanner module, with LED tripod attached to it, and the universal microscope part (identical for all of the microscope versions). Most of the parts and optics hold sufficiently well with pressure fit. In rare occasions it is necessary to add some glue, as the tolerances of 3D printing and different parts are not good enough to ensure a stable press fit in all cases. **d**, MEMS scanner module. First the collimation lens assembly is inserted, then the scan lens assembly, then the MEMS scanner. In our experience fitting the MEMS scanner in its socket is sufficient and no further alignment is necessary. **e**, universal microscope part. The dichroic mirror is first inserted from the side. Then the tube lens is positioned through its aperture. The condenser lens is added from top. The objective lens can be mounted and dismounted anytime and is usually added just before an imaging session. See also Supplementary movie 7.

**Supplementary Figure 11.**
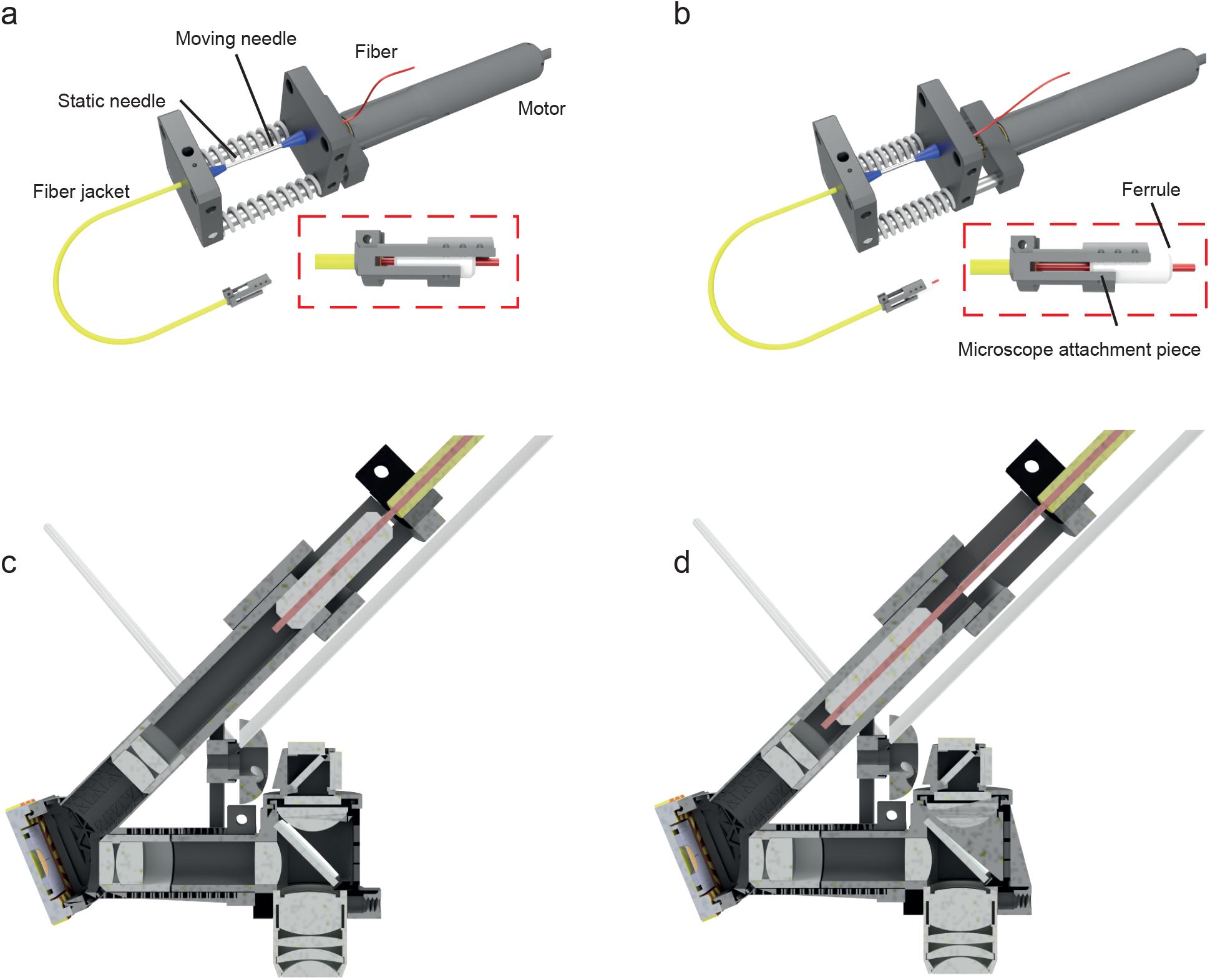
Implementation of the motorized mechanical fiber tip shift actuated from the laser table side of the fiber. **a**, motorized optical table setup for actuating the fiber tip, with the fiber tip, glued inside a ceramic ferrule, in its most retracted position. The inlay shows an enlarged view of the ferrule and microscope attachment piece. **b**, as in a, but with fiber tip in its most extended position. **c**, cross-section of the microscope with the fiber passively sliding in a cylindrical sleeve, show in its most retracted position. d, as in c, but with the fiber tip in its most extended position. See also Supplementary movie 6.

**Supplementary Movie 1. Z-stacks acquired using either the head-mounted z-drive or with the miniature microscope mounted on an external micromanipulator.** Z-stacks from the brain surface to the white matter. All data has been acquired with high Z-range version of the microscope in primary visual cortex of an anesthetized mouse. Images are overlays of jGCaMP7f fluorescence (green) and the third harmonic signal (blue). From left to right: side projection of the stack acquired with the headmounted z-drive; image stack acquired with the head-mounted z-drive; image stack acquired with the miniature microscope mounted on an external micromanipulator; side-projection of the z-stack acquired using the external micromanipulator. Note that the first side projection has been scaled approximately by comparing visible structures with the conventional z-drive micromanipulator.

**Supplementary Movie 2. Comparison of raw imaging data, a 3 frame rolling average and de-noised imaging data.** All data has been acquired with the high Z-range version of the microscope in visual cortex of a freely moving mouse. Movies show green fluorescence from jGCaMP7f. Imaging is shown in real time. Top row, from left to right: raw data; 3-frame rolling average, imaging data processed with DeepCAD (Li, Zhang et al. 2021). Bottom row: fluorescence traces calculated from the imaging data in the same column from the neurons indicated by the colored circles in the movies.

**Supplementary Movie 3. Imaging and mouse behavior with lit and dark environment.** All data has been acquired with high Z-range version of the microscope in visual cortex of a freely moving mouse. Movies show green fluorescence from jGCaMP7f. Imaging is shown in real time. Top left: imaging data processed with DeepCAD (Li, Zhang et al. 2021). Top right: visible spectrum over-head camera view with outer front (white), outer left (red) and outer right (green) infra-red LED positions labelled. Bottom: fluorescence traces from the neurons indicated in the fluorescence images. Scale bar applies to all traces.

**Supplementary Movie 4. Remote depth navigation between cortical layers 4 and 6.** All data has been acquired with high Z-range version of the microscope in visual cortex of a freely moving mouse. Fluorescence from jGCaMP7f shown in green, third harmonic signal in blue. Imaging is shown three times real speed. Left: imaging data shown with a 10 frame rolling average, while the head-mounted z-drive is used to change the imaging depth from layer 6 to the bottom of the cortex, with white matter visible in the blue channel, then up to layer 4. At layer 4 the field of view was also shifted by applying voltage offset to the MEMS scanner, adjusting the lateral position of the field of view. Right: visible spectrum over-head camera view, with outer front (white), outer left (red) and outer right (green) infrared LEDs positions labelled.

**Supplementary Movie 5. Animated CAD models showing the procedure for mounting the microscope on the head plate.** Animation illustrating the sequential steps during the positioning and mounting procedure and for an imaging experiment. The individual steps are shown in Supplementary figure 8.

**Supplementary Movie 6. Animated CAD models showing the mechanical Z-Drive setup.** Animation of operation of the mechanical z-drive as shown in Supplementary figure 11.

**Supplementary Movie 7. Animated CAD models showing the process of assembling the microscope.** Animation of the assembly process as described in Supplementary figure 10.

